# LAS3R: A simple, secure, scalable, and robust framework for deploying lab automation devices

**DOI:** 10.64898/2026.04.08.716564

**Authors:** Kavi Haria Shah, Gos Micklem

## Abstract

Laboratory automation can greatly accelerate experiments and data collection, yet building automated systems often requires substantial programming and electronics expertise, and few frameworks are targeted at deploying many devices. We present LAS3R, a low-cost, open-source framework that enables researchers with minimal technical expertise to rapidly prototype, deploy, remotely control, and collect data from multiple custom-built laboratory devices while maintaining strong security and reliability throughout the process—from early prototyping to routine operation. The system is built around a central hub to which multiple lab devices connect. This hub can be set up on a Raspberry Pi (a small, low-cost single-board computer) in under fifteen minutes. In the setup process, code is automatically generated for ESP32 microcontroller boards that control the hardware. Users can choose from a list of preconfigured ESP32 devices, for example a bioreactor, or use a template that provides base code for many common automation tasks, which they can then easily customise using the beginner-friendly Arduino platform. The ESP32 devices connect through a secure Wi-Fi network hosted by the Raspberry Pi that encrypts communication, and ensures only authorised hardware can join, helping safeguard experimental data and institutional networks, even while prototyping. We demonstrate the framework with two applications—a turbidostat bioreactor and a light-level controller—and show that it can simultaneously manage eight devices with 24 sensors. Robustness was evaluated through single-point-of-failure analysis, confirming continued operation during mains power or network interruptions. Comprehensive documentation, aimed at wet lab researchers, enables users to understand, build, and adapt the system, making it both a practical laboratory automation platform, including for those in low-resource settings, and a teaching resource. This paper is intended to be a technical evaluation of the architecture. Those wishing to deploy the system should refer to the online documentation at kavihshah.github.io/LAS3R.

## Introduction

Expanding lab automation promises to increase experiment throughput, improve reproducibility, and enable continuous monitoring of environmental conditions, while reducing manual effort, thereby freeing researchers to focus on other tasks. However, greater adoption is limited by the high cost of commercial automation platforms, their inflexibility for novel or custom applications, and the technical expertise required to design, manage, and operate such systems. Particularly in resource-constrained settings, the costs of commercial automation systems are prohibitive [1]. Standardisation and cost reduction have therefore been identified as key factors to increase accessibility [2].

In response to the limited accessibility and restricted flexibility of commercial solutions, there has been growing interest in low-cost, open-source laboratory devices. Recent examples include bioreactors such as Chi.Bio [3]; optical analysis instruments and imaging platforms like the qByte [4], FluoroPi [5], and OpenFlexure Microscope [6]; and liquid-handling systems such as the OpenLH [7], to name just a few. These community-driven tools enable laboratories to prototype, customize and use automation solutions tailored to specific experimental needs, accelerating research while reducing dependence on expensive commercial equipment. Despite their success, challenges remain: varying levels of documentation, differences in device setup and limited standardisation, and the technical expertise required to modify some devices hinder broader adoption. Moreover, few systems offer integrated frameworks for the simultaneous development, control, and monitoring of multiple instruments, with many relying on ad hoc communication protocols which vary from HTTP to serial interfaces, limiting scalability, flexibility, and security. The requirements for a fully functional lab automation framework are outlined in Table 1.

**Table 1.**
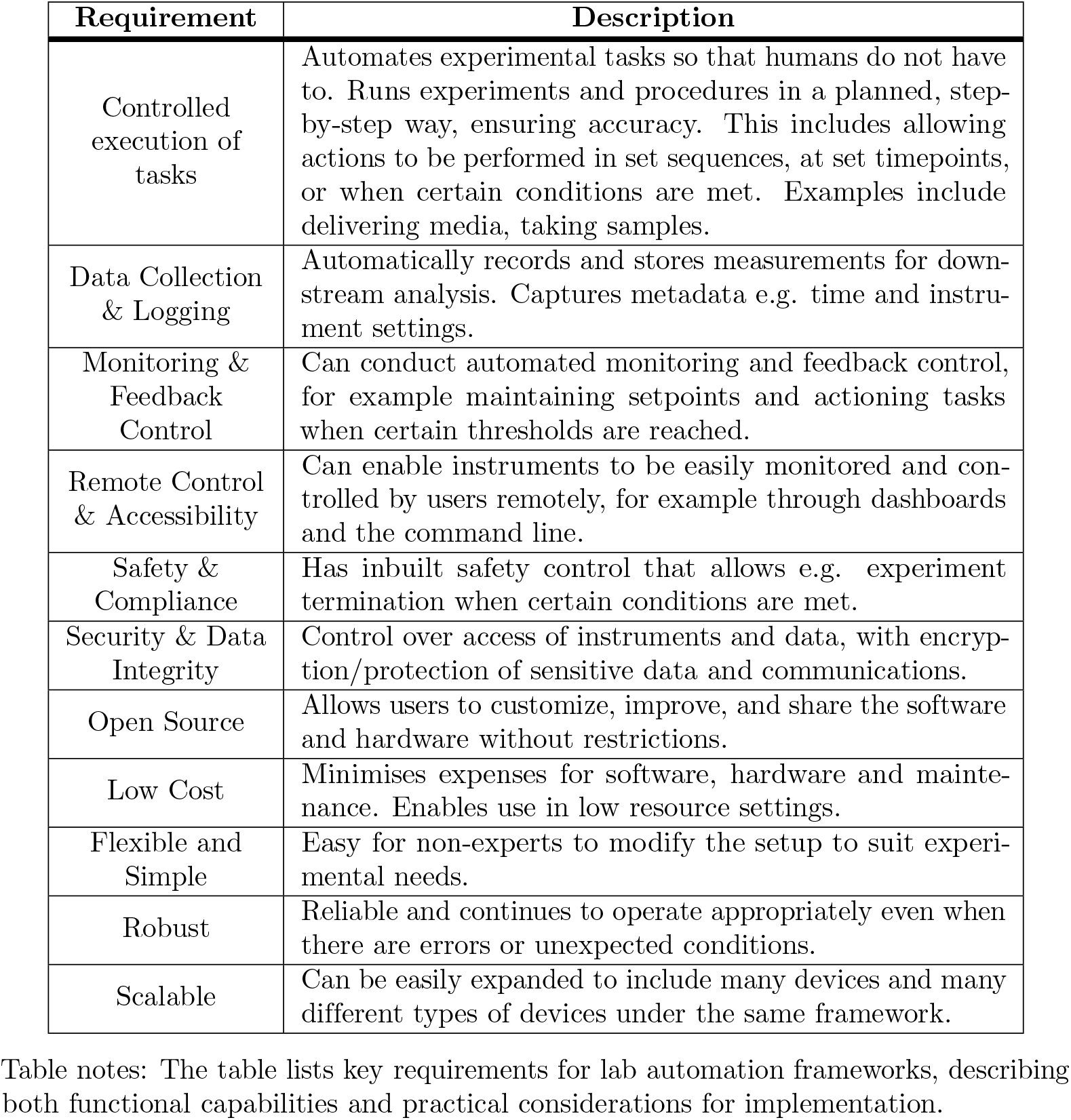
Requirements for a lab automation framework. Table notes: The table lists key requirements for lab automation frameworks, describing both functional capabilities and practical considerations for implementation.

| Requirement | Description |
| --- | --- |
| Controlled execution of tasks | Automates experimental tasks so that humans do not have to. Runs experiments and procedures in a planned, step-by-step way, ensuring accuracy. This includes allowing actions to be performed in set sequences, at set timepoints, or when certain conditions are met. Examples include delivering media, taking samples. |
| Data Collection & Logging | Automatically records and stores measurements for downstream analysis. Captures metadata e.g. time and instrument settings. |
| Monitoring & Feedback Control | Can conduct automated monitoring and feedback control, for example maintaining setpoints and actioning tasks when certain thresholds are reached. |
| Remote Control & Accessibility | Can enable instruments to be easily monitored and controlled by users remotely, for example through dashboards and the command line. |
| Safety & Compliance | Has inbuilt safety control that allows e.g. experiment termination when certain conditions are met. |
| Security & Data Integrity | Control over access of instruments and data, with encryption/protection of sensitive data and communications. |
| Open Source | Allows users to customize, improve, and share the software and hardware without restrictions. |
| Low Cost | Minimises expenses for software, hardware and maintenance. Enables use in low resource settings. |
| Flexible and Simple | Easy for non-experts to modify the setup to suit experimental needs. |
| Robust | Reliable and continues to operate appropriately even when there are errors or unexpected conditions. |
| Scalable | Can be easily expanded to include many devices and many different types of devices under the same framework. |

One notable effort to address these limitations is LabThings, an open-source implementation of the W3C Web Things architecture [8]. LabThings allows instruments to be automatically discovered over standard IP networks, describing their functionality via Thing Descriptions, and enabling interaction and automation through HTTP and WebSocket APIs. Existing lab instruments can be adapted for LabThings using low-cost intermediary devices such as the Raspberry Pi or Arduino. By providing a standardised, interoperable framework for experiment planning, resource management, asynchronous execution, and data handling, LabThings enables laboratories to move away from costly, proprietary solutions and adopt sustainable, accessible, and modernised lab automation practices.

However, LabThings presents a relatively high technical barrier for users with a primarily biological background, and focuses on device interfacing rather than serving as a base for prototyping novel automation devices. Furthermore, its reliance on HTTP/HTTPS can introduce additional computational and protocol overhead compared to other lightweight communication protocols, particularly in scenarios involving very frequent, small payload messages common in data streaming and collection. As the number of interfaced devices scales, especially in architectures with multiple data consumers, this can increase network and processing load, potentially impacting system performance and requiring more complex architectures to manage, particularly on resource-constrained hardware.

We present a simpler, robust, and scalable framework for securely prototyping novel devices, targeted at biologists with limited coding experience, and using the more efficient MQTT protocol instead of HTTP/HTTPS for communication and data collection. MQTT provides a simple and scalable publisher-subscriber model managed by a broker, and also has built-in quality of service (QoS) levels to manage message delivery reliability.

A further important consideration is the security of self-built devices on institutional networks. Academic and public institutions are increasingly at risk of cyberattacks and frequently handle sensitive data [9]. Despite this, many lack a dedicated isolated prototyping network, or a separate network specifically configured for enhanced security. As a result, all traffic, including low and high security operations and prototyping activities, shares the same network. Furthermore, even though commercial systems are not always significantly more secure, institutional policies often limit open-source hardware development, deployment, and remote control. LabThings, built on Flask (LabThings GitHub) [8], demonstrates this limitation: its example server scripts use plain HTTP on the standard development port (run via app.run(host=“0.0.0.0”,port=5000)) and do not include SSL/TLS (Secure Sockets Layer/Transport Layer Security) configuration by default, meaning the server communicates over unencrypted HTTP unless explicitly configured. This default behavior poses security concerns, particularly when connecting multiple devices or prototyping within institutional networks, highlighting the need for frameworks that balance scalability, usability, and robust security. Our framework addresses this concern by enforcing security standards by default, including TLS (Transport Layer Security) for encrypted communications and certificate-based authentication, so users can focus on functionality without compromising security, balancing protection with usability and development flexibility.

## Materials and methods

### Software and hardware framework

The developed framework was designed to use a Linux machine with Access Point Wi-Fi capability as a central node, and was evaluated primarily on Raspberry Pi 4 and Raspberry Pi 5 devices [10] running Raspberry Pi OS 64-bit (Debian-based, release 13 May 2025 [11], imaged via Raspberry Pi Imager v1.8.5 [12]). In the default setup used in the subsequent experiments, all core services described below were hosted or run from this single Unix machine. Local public key infrastructure(PKI) was implemented with OpenSSL [13] enabling certificate-based authentication. An isolated local network was hosted using the network services hostapd [14] (access point control), dnsmasq [15] (DNS/IP assignment), and FreeRADIUS [16] (WPA-Enterprise authentication). Encrypted communication via MQTT over TLS was facilitated by the Eclipse Mosquitto broker [17]. Certificate Revocation Lists (CRLs) were hosted on an Apache HTTP Server [18]. Host-level security employed UFW [19] and sshd with key-based access control. Data acquisition and visualization used Telegraf [20] and InfluxDB v2 [21]. Control and automation scripts were developed in Python, employing the Paho-MQTT library [22] to interface with the MQTT broker. Background services and startup configurations were managed through standard Linux daemon controllers. The setup scripts written in Bash are designed to run from the central node, and are available at the github repository: github.com/KaviHShah/LAS3R.

Microcontroller integration used the Arduino IDE [23] with the ESP32 platform (Espressif Systems, board package v3.1.1 [24]). Firmware was written in arduino C++ utilising the WiFi101 (Arduino v0.16.1) [25], WiFiClientSecure [26], esp eap client [27], and PubSubClient(v2.8) [28] libraries. The DFRobot Beetle ESP32 (DFR0575) [29] and Adafruit HUZZAH32 ESP32 Feather Board [30] were the tested microcontrollers. Microcontroller code for various devices as well as template code is generated in an automated fashion during setup, and is available at the github repository: github.com/KaviHShah/LAS3R, and can be pushed to ESP32 devices using the Arduino IDE. In addition, in depth documentation and a quick start user guide is available at kavihshah.github.io/LAS3R.

### Bioreactor setup and experiments

The turbidostat bioreactor was constructed by augmenting a heating mantle with inbuilt temperature control. Detailed hardware specifications are available on the project repository (github.com/KaviHShah/LAS3R Bioreactor). Briefly, an inline optical density (OD) sensor was implemented using an OPT101 photodiode coupled with a Thorlabs high power LED (wavelength: 600nm LED600L, 600 nm LED with a Glass Lens, 3 mW, TO-18), and integrated via a peristaltic pump (GROTHEN 1*3 mm 12V DC) to allow continuous sampling of the culture. Influx and efflux peristaltic pumps (GROTHEN 1*3 mm, 3*5 mm 12V DC) were used to feed in fresh media and remove waste thereby allowing control of the turbidity of the culture, for example maintenance at a set point.

For the initial growth curve experiment, 50 mL of E. coli MG1655 was cultured overnight in the bioreactor at 37 °C in LB medium without addition of fresh media. The OD sensor was used to monitor growth.

To determine an accurate growth rate, within the turbidostat bioreactor a 50 mL MG1655 culture was diluted to an initial OD_600_ of 0.55 a.u. (corresponding to light sensor value 1800 a.u.) using fresh LB media and then allowed to grow until the culture reached OD_600_ ≈ 0.60 a.u. (light sensor value 1900 a.u.) before being diluted again. This regime was conducted repeatedly to generate replicates and the average growth rate calculated. Control of set points and thresholds was performed using a Python script running on a Raspberry Pi, with communication facilitated via MQTT. This script is available on the github repository github.com/KaviHShah/LAS3R (mqtt publish setup script). The turbidostat OD measurements were calibrated against a spectrophotometer (GeneQuant 1300 Spectrophotometer, Biochrom Ltd; 1 mL plastic cuvette with path length 10 mm) to generate the experiment calibration curve and retrieve OD values.

### Light-controller prototyping

The template code provided during system setup was used to rapidly prototype an LED light-intensity controller within two hours. The controller maintains prescribed illumination levels by regulating LED (Thorlabs wavelength: 600nm LED600L, 600 nm LED with a Glass Lens, 3 mW, TO-18) outputs through pulse-width modulation (PWM). The original Δ*t*-aware PID-control template was implemented with appropriate integral windup protection through cumulative error clamping, and actuator saturation. Output rate limiting following setpoint changes was also added to improve stability. PID parameters were tuned using a combination of the Ziegler–Nichols method and subsequent manual refinement. Additional functionality was implemented to enable automatic failover in the event of system faults. A Python script was executed by a system daemon to periodically reassert setpoint values (see mqtt publish setup script within github.com/KaviHShah/LAS3R), and the hostapd setup modified to support rapid reconnection of devices. For details see the hostapd setup script within github.com/KaviHShah/LAS3R.

### Robustness testing utilising the light-controller

The prototyped light-controller device was evaluated through a series of single-point-of-failure (SPOF) tests to assess system robustness. During all tests, a Python script using the Paho MQTT client library published control signals to maintain the LED output at a target intensity of 50 arbitrary units (as defined by the photodiode). Each failure mode perturbation was introduced at 2 min and was repeated three times sequentially with a period of 5 minutes. Where specified the perturbation was resolved after a minute.

#### Power-related failure modes

Power was stopped at 2 min and resumed after a one minute break.

- Loss and restoration of power to the Raspberry Pi without an uninterruptible power supply (UPS).
- Loss and restoration of power to the ESP32 without a UPS.
- Loss and restoration of power to the Raspberry Pi with a UPS.
- Loss and restoration of power to the ESP32 with a backup battery power supply.

### Service-level failure modes (network, communication and data-collection)

- hostapd service interruption: systemctl stop hostapd.service was issued at 2 minutes and the service was restored at 3 minutes.
- Mosquitto MQTT broker interruption: systemctl stop mosquitto.service and subsequent restart at 3 minutes.
- Forced termination (SIGKILL) of networking services with automatic recovery by service daemons:
  - kill -9 on hostapd
  - kill -9 on mosquitto
  - kill -9 on telegraf
  - kill -9 on influxdb
  - kill -9 on freeradius

#### Physical network connectivity failure modes

- Loss of Wi-Fi connectivity by physically exiting and re-entering the access point range. The device was moved out of range at 2 minutes and re-entered at 3 minutes.

#### Hardware-level failure modes

- Simulated LED failure: One LED was disconnected during operation to trigger the controller’s automatic switch to a backup LED. A 10-second delay was incorporated into the ESP32 firmware to facilitate visual confirmation of the handover.

#### Fail safe behaviour

- Loss of Wi-Fi connectivity with ESP32 fail-safe behaviour enabled: In this configuration, the ESP32 script was configured to disable the LEDs when external control or Wi-Fi was unavailable.

### Scalability testing

Data collection and control were evaluated using eight simultaneously connected ESP3232 devices, each simulating three sensors (24 sensors total), with continuous data streaming at rates exceeding 1 reading per second per sensor. In the simulation each sensor returns the set value unique to the device + a random error term to simulate a real sensor. The device set points were set at 20 (a.u.) intervals apart for visualisation clarity. A mixture of Addafruit ESP32 feather (HUZZAH32) and Dfrobot ESP32 beetle (DFR0575) devices were used.

### Code and documentation

All code required in order to rapidly set up the framework is in the Github repository: github.com/KaviHShah/LAS3R. In addition we provide detailed documentation and tutorials on the website kavihshah.github.io/LAS3R.

## Results

### Framework Overview

At its core, the system utilises a secure WPA-Enterprise local Wi-Fi network, hosted on a central Linux-based machine with a suitable Wi-Fi chip, to which Wi-Fi–enabled microcontrollers like ESP32s can connect (Fig 1). A Raspberry Pi was used as a low-cost central Linux node during development, demonstrating the minimal hardware requirements for the setup.

**Fig 1.**
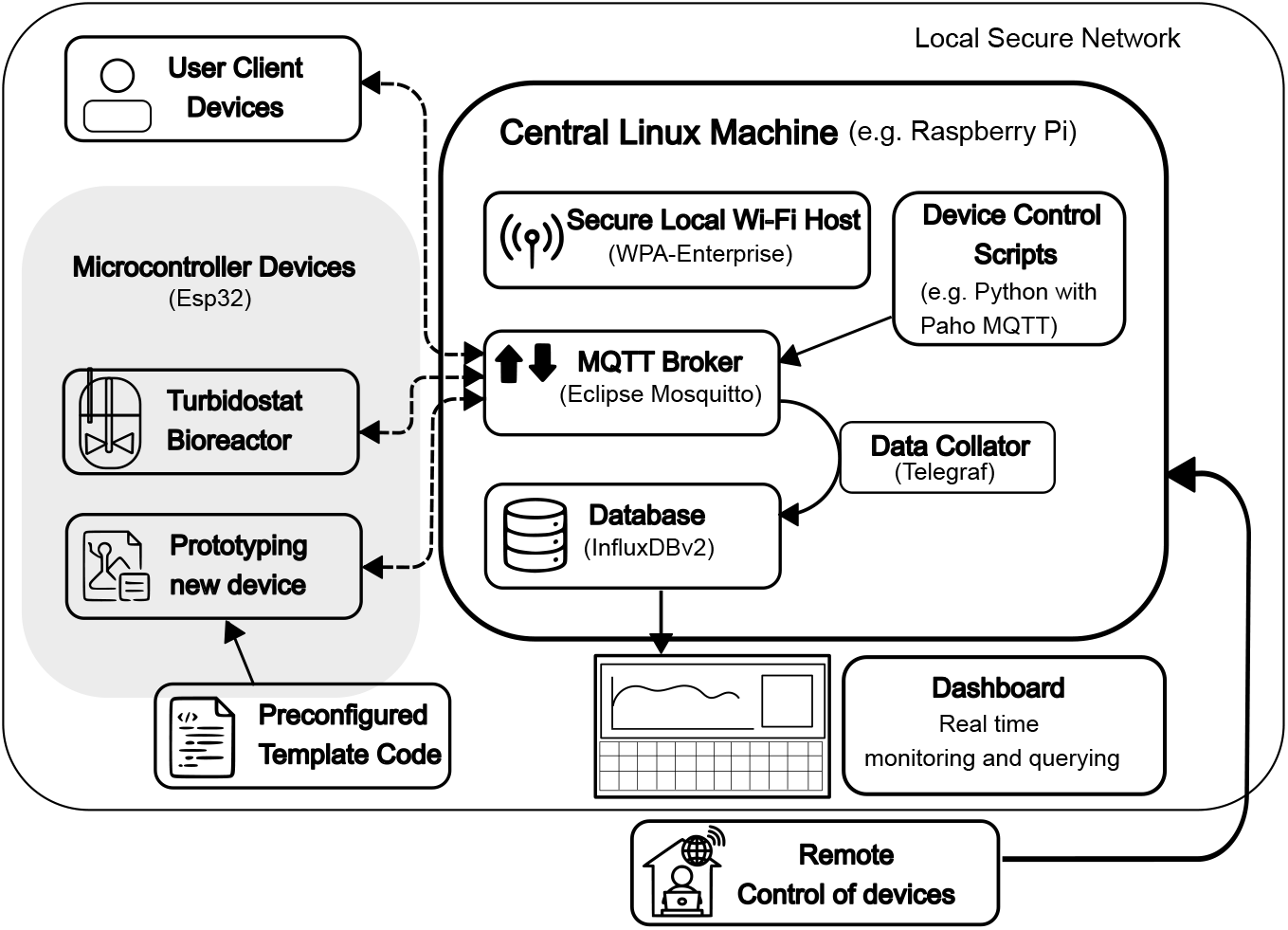
Overview of the Lab Automation Framework. A central linux machine (Raspberry Pi) is used to host a secure local Wi-Fi network. In addition, the central machine hosts an MQTT Broker for communication, and database (InfluxDB v2) for data storage. Microcontroller devices, as well as user devices, can connect to the local network. During setup, preconfigured device code can be generated (e.g. turbidostat bioreactor), or alternatively template code can be generated, which can then be used to rapidly prototype new devices. MQTT scripts run on the central machine, or MQTT commands by users on the network, are used to control devices externally. Remote control of devices from outside of the local network can only be done through SSH into the central machine. Device-generated data are published on MQTT channels, collated by the Data Collator (Telegraf), and stored in the database. InfluxDB provides a dashboard for real-time monitoring and data querying.

Device control, communication, and data streaming, are implemented through the MQTT protocol utilising the Eclipse Mosquitto broker [17], while experimental data is collected and stored in an InfluxDBv2 database, all hosted on the central linux machine. During setup, preconfigured ESP32 firmware for user-specified devices—including a turbidostat bioreactor—or alternatively template code for the prototyping of new devices is automatically generated (Fig 1). This template code enables secure access to the network during prototyping, and covers most automation and communication needs, requiring only minimal modification on the beginner-friendly Arduino IDE programming platform to produce novel devices. The entire framework, including microcontroller code generation, can be set up on a Raspberry Pi in under fifteen minutes. The setup scripts automatically handle user account creation and other necessary utilities as part of this process. For technical details, please refer to the methods section and setup scripts, as well as the detailed documentation on the website.

### Framework security

The framework establishes a secure environment for device prototyping and deployment through an isolated local network (Fig 2A). Network access is restricted to authorised devices and users, which are required to present valid certificates (WPA-Enterprise) [31] issued by a local Public Key Infrastructure (PKI). All communication within the network is protected using Transport Layer Security (TLS) [32] protocols also underpinned by certificates, ensuring data confidentiality through encryption and integrity during transmission. This combination of certificate-based authentication and encrypted communication ensures robust protection against unauthorized access and eavesdropping (Fig 2B).

**Fig 2.**
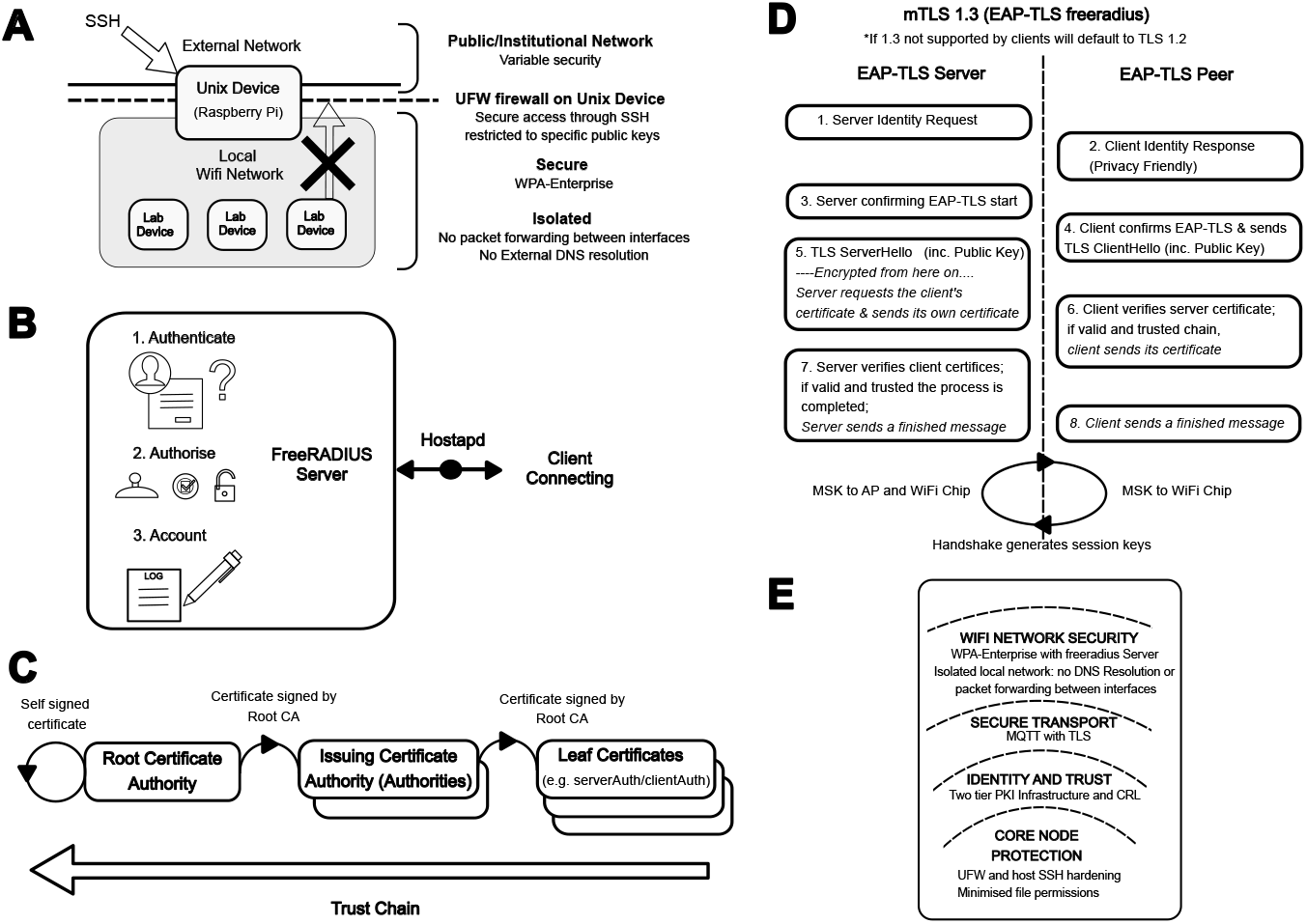
Framework Security. **A:** A Unix-based device (Raspberry Pi) is used to host a local network. The network is secured using WPA-Enterprise, requiring certificate-based authentication for laboratory devices to connect. Network isolation is enforced by disabling DNS resolution and packet forwarding between interfaces, preventing connected devices from accessing the wider internet. The central machine is further secured using UFW (Uncomplicated Firewall), with SSH access restricted to authorised public keys when accessed via the external institutional network. **B:** When a client connects to the secure network, hostapd forwards the authentication request to FreeRADIUS for validation. FreeRADIUS performs certificate-based authentication, authorizes access by verifying credentials and checking against the certificate revocation list (CRL), and logs connection events. **C:** A two-tiered Public Key Infrastructure (PKI) is employed. The root certificate authority (CA) resides at the top of the trust chain and is self-signed. One or more issuing certificate authorities, whose certificates are signed by the root CA, are responsible for issuing and signing end-entity (leaf) certificates. **D:** A high-level overview of mutual TLS (mTLS) 1.3 authentication implemented using EAP-TLS [31] (Extensible Authentication Protocol–Transport Layer Security). Both the peer (connecting client) and server verify each other. If TLS 1.3 is not supported by clients, the current implementation allows FreeRADIUS to fall back to TLS 1.2. The process begins with the client initiating a connection. In response to a request from the authenticator access point (1), the client provides an initial identity (2). This identity may be anonymized (e.g., an outer identity) since the exchange occurs prior to establishment of the encrypted TLS tunnel. (3) The server (via RADIUS) selects and initiates the EAP-TLS method. (4) A TLS handshake is then performed within the EAP exchange, beginning with the client sending a ClientHello, which includes supported cryptographic parameters and key share information. (5) The server responds with a ServerHello, containing its cryptographic parameters and public key share. Following this, the server sends its certificate and, if required, a CertificateRequest for the client certificate. In TLS 1.3, all handshake messages after the ServerHello—including the server certificate and certificate request—are encrypted and transmitted together in a single flight. (6) The client validates the server’s certificate against a trusted certificate authority. If verification succeeds, the client sends its own certificate and proves possession of the corresponding private key. (7) The server validates the client certificate, including revocation checks, completing mutual authentication. (8) Both parties exchange Finished messages to confirm successful establishment of the TLS session. Upon successful completion, keying material (the Master Session Key, MSK) is derived from the TLS session and exported to the authenticator (access point). The access point and client then use this material to derive transient session keys (e.g., via the 4-way handshake in WPA2/WPA3-Enterprise) to secure subsequent wireless communication. **E:** Summary of the security measures in place in the automation framework.

#### Certificate management

A two-tiered local Public Key Infrastructure (PKI) is employed, consisting of a root Certificate Authority (CA) and an intermediate CA (Fig 2C). The intermediate CA is responsible for generating device and user certificates using OpenSSL [13]. This hierarchical structure provides a verifiable trust chain, enabling secure authentication and authorization of network participants. Certificates are mandatory for joining the local network and for TLS communication, effectively preventing untrusted devices from accessing network resources.

#### Network security and communication

The local network is implemented on a Linux host using hostapd [14] to provide a Wi-Fi access point, FreeRADIUS [16] for WPA-Enterprise authentication, and Dnsmasq [15] to handle DHCP and internal IP assignment. The local network operates without external DNS (Domain Name System) resolution and packet forwarding between interfaces, relying exclusively on internal IP addressing (Fig 2A). This increases security by preventing malware, misconfigured devices, or rogue software from sending data outside the network. Similarly, devices connected solely to the network cannot access the wider internet. Network authentication is implemented using WPA-Enterprise–grade (Wi-Fi Protected Access) using EAP-TLS [31](Extensible Authentication Protocol–Transport Layer Security) protocols via FreeRADIUS (Fig 2D), with enforcement of Certificate Revocation Lists (CRLs) to prevent access by compromised or revoked credentials. Device-to-device communication is secured using TLS-protected MQTT, ensuring confidentiality and integrity of transmitted data.

#### Host-level security and access control

The network host serves as the sole access point for users without network certificates and as the primary interface for configuration and setup modifications. Host-level security is enforced through a combination of custom firewall rules implemented via UFW [19] (Uncomplicated Firewall), secure shell (SSHD) configurations, and strict user management policies. SSH access is restricted to a designated entry node, such as a Raspberry Pi, with key-pair authentication required to ensure that only authorized personnel can connect (Fig 2A). File system permissions are carefully controlled, with administrative privileges regulating access to sensitive configuration files and certificates. Collectively, these measures provide robust authentication, granular access control, and protection of sensitive data throughout the prototyping and deployment process. While not part of the default configuration, disk encryption is recommended, particularly for setups with easily accessible hardware or extended periods of downtime.

#### Balancing usability and security

For environments handling highly sensitive data, it is advisable to follow the documentation in detail and implement the framework manually with a security-first approach. Recommended practices include, but are not limited to, generating the PKI infrastructure on an air-gapped system, utilising Hardware Security Modules (HSMs), pre-installing and verifying all required packages, and performing offline configuration. Additionally, selecting the latest encryption standards (e.g., elliptic curve cryptography), minimising certificate validity periods, enforcing stricter certificate management policies, and mandating the use of TLS 1.3, are strongly encouraged. Modifications to microcontroller scripts should also be carried out as necessary to align with these security measures. Further guidance on these procedures is provided in the documentation, though implementing them requires a certain level of technical expertise. The setup script achieves a practical balance between usability and security, rendering it suitable for the majority of academic contexts, and providing markedly higher security compared with common current practices.

#### Summary of security measures

- Isolated local network requiring PKI-issued certificates for access.
- Two-tier PKI (root CA and intermediate CA) for certificate generation and trust management.
- WPA-Enterprise authentication via FreeRADIUS with CRL enforcement.
- TLS-protected MQTT for secure communication between devices.
- Host-level security through UFW, SSHD rules, and controlled user permissions.
- SSH access limited to a single entry node with key-pair authentication.
- File permissions managed access restricted to administrators to protect sensitive data.

These combined measures create a robust security framework suitable for secure device prototyping and deployment (Fig 2E). While further modifications to the setup can be implemented for high security contexts (as detailed above and in the website documentation pages), for most laboratory applications, the setup, combined with adherence to good practices, offers an effective balance between good security and usability.

### Framework Communication

Communication, control, and data collection within the LAS3R framework are facilitated via MQTT over TLS (Fig 3A), utilising the Eclipse Mosquitto broker [17]. MQTT (Message Queuing Telemetry Transport) is a lightweight, publish/subscribe messaging protocol designed for efficient, reliable exchange of small, frequent payloads, making it particularly suitable for laboratory automation environments. Unlike HTTP, MQTT is optimised for scenarios requiring frequent bidirectional communication between multiple devices [33]. While MQTT is typically optimised for compact data transmission, it supports arbitrary binary payloads, and does enable the transfer of diverse data types, including images, if necessary (some template HTTP code is also provided which may be optimal for certain data type transfer use cases). The LAS3R framework performs reliably under constrained bandwidth and limited processing power, and offers MQTT over TLS for security. Communication is coordinated through a central broker, which ensures dependable message delivery between publishers and subscribers (Fig 3A). Data is organized into topics, with publishers transmitting to specific topics and subscribers receiving relevant updates. Persistent TCP connections, combined with configurable Quality of Service (QoS) levels, ensure consistent and reliable message transmission.

**Fig 3.**
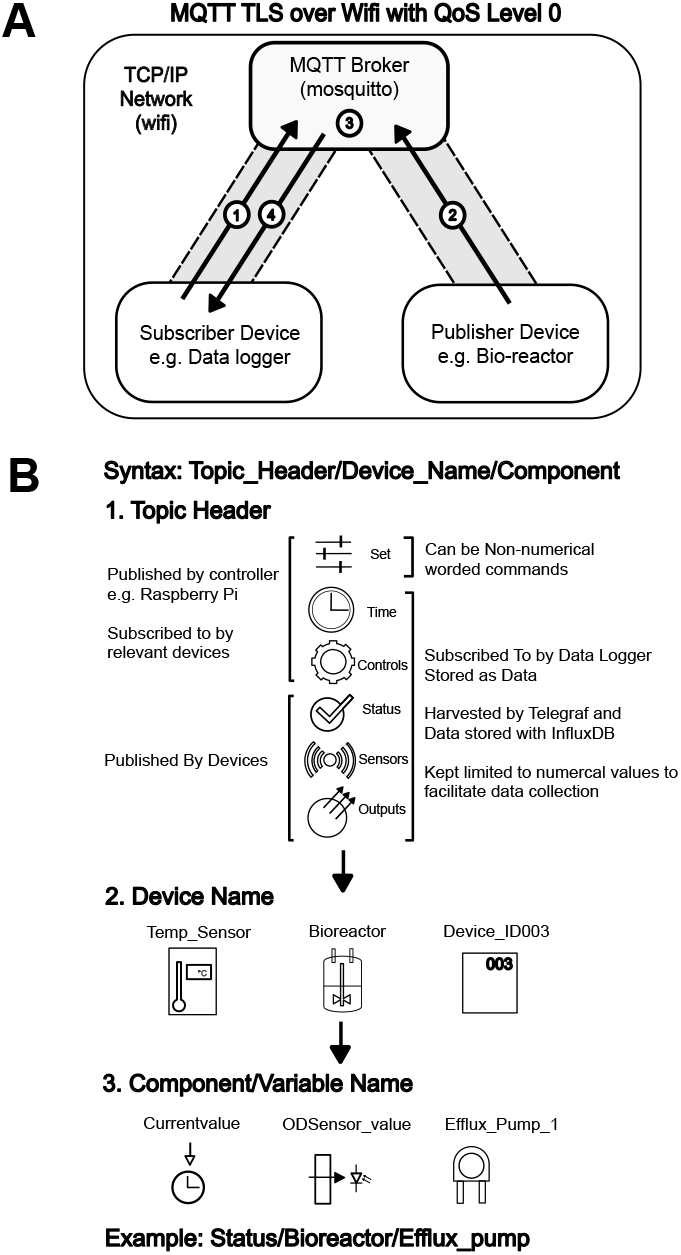
Communication overview. **A:** MQTT operates over a TCP/IP network — in our case, over a Wi-Fi network — and follows a publish-subscribe communication model. The MQTT broker is responsible for forwarding messages received from a publisher to all clients subscribed to the topic (channel) the message was published to. TLS is used in our setup, meaning that clients and the broker authenticate each other using certificates, and all messages are encrypted in transit. If a subscriber is not connected at the time a message is published, the message is lost. Clients, which can be publishers, subscribers, or both, connect to the MQTT broker over a long-lived, bidirectional TLS-encrypted TCP connection (grey band). (1) Subscribers subscribe to topics. For Quality of Service Level (QoS) 0, this subscription must occur before any messages are published to that topic, otherwise, messages will not be received as the broker does not store messages, nor does it require acknowledgement from subscribers (2) Publishing devices (e.g. a bioreactor) publish sensor data to a topic. (3) The broker receives the message and forwards it to all clients subscribed to that topic. (4) Subscribed clients receive the message in real time. **B:** Framework syntax for MQTT channels consisting of three layers. In the top layer, there are six key headers reflecting the type of content. The Set, Time and Controls channel headers are published by controllers and subscribed to by relevant devices, whereas the status, sensors and output channel headers are used by the devices to publish. All channels apart from the set channel are collected as data and kept limited to numerical content for ease of data collection and analysis. The second layer consists of the device name e.g. Temp Sensor or Bioreactor; whereas the third and final layer consists of the component or variable name.

By leveraging MQTT over a Wi-Fi network instead of physical connections such as serial communication, the LAs3R framework simplifies wiring, enhances scalability, allows efficient management of multiple devices, and even supports the creation of mobile devices. The LAS3R automation framework further defines a default set of communication channels to interface with data collection (Fig 3B). Standard topic names follow the format: Topic Header/Device Name/Component, with the following topic headers: Set, Time, Controls, Status, Sensors, and Outputs. The Set topic is reserved for device control commands and is not logged by the data collector, whereas all other topic headers are logged by default and stored in the database. Users may customize this topic structure to suit specific experimental or deployment requirements.

### Data Storage, Visualisation and Querying

Data collection and storage were managed using InfluxDB v2 [21], an open-source time-series database developed by InfluxData. InfluxDB is designed for efficient storage and analysis of high-volume, time-stamped data and supports querying the data and real time visualisation with a provided GUI (Fig 4A). Its schema-less architecture allows data to be ingested with minimal preconfiguration, making it highly adaptable to dynamic experimental environments. The LAS3R framework uses InfluxDB interfaced with the associated Telegraf [20] data collection tool to harvest data from select MQTT channels. InfluxDB version 2, distributed under the permissive open source MIT license, is a mature and widely adopted solution that fully met our requirements and is likely to satisfy the needs of most academic biology laboratories handling time-series data. Alternatives are discussed further in the online documentation (kavihshah.github.io/LAS3R). Additionally code for the ESP32 devices to create dashboard web-interfaces is included within the template code generated at setup (github.com/KaviHShah/LAS3R, Fig 4B).

**Fig 4.**
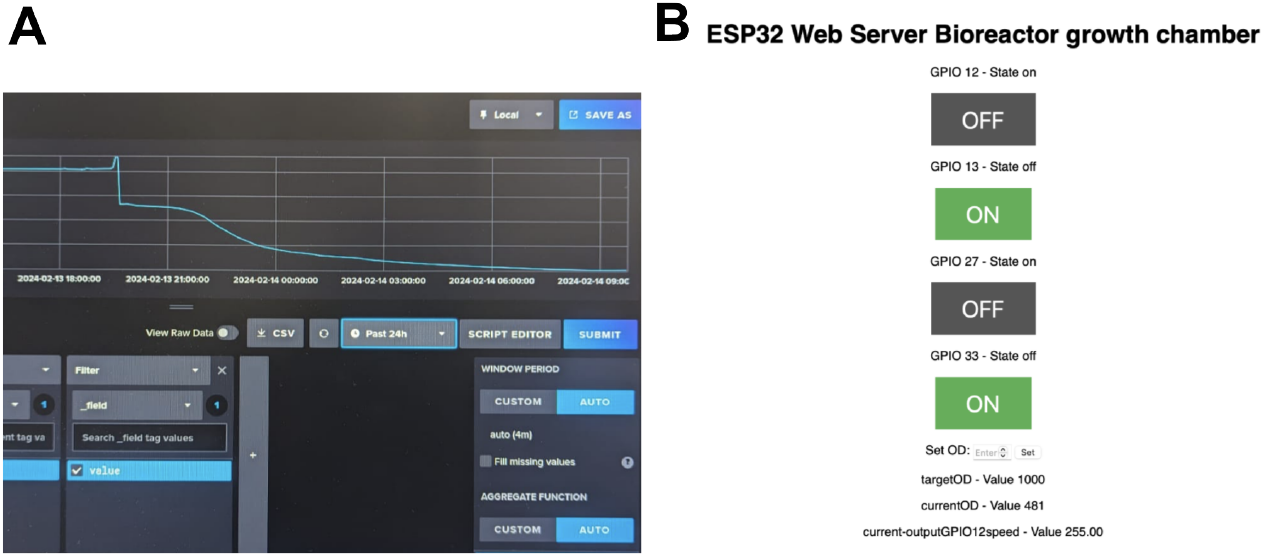
Data Visualisation. **A:** The InfluxDBv2 graphical user interface for visualising and querying data. **B:** A web server hosted by the ESP32 device, in this case the bioreactor, that can be used to monitor device status and control the device remotely (Code provided in the template)

### Framework Ease of Use

The LAS3R framework has been developed to ensure that the user experience, particularly for non-experts, is as straightforward and streamlined as possible. An in-depth understanding of all underlying tools is not required to use, adapt, or prototype devices with the framework. To facilitate this, the documentation includes a Quick Start Guide that leads users through the initial setup, and extensive documentation provides in-depth guidance for modifying the framework.

The central component of the LAS3R framework is the ‘setup.sh’ script, which is the only script that must be executed. Upon execution, the script prompts users for all required inputs; users simply provide responses, guided by the Quick Start Guide. The ‘setup.sh’ script subsequently invokes all the individual setup scripts necessary for the system configuration. Within this setup script, users can select which sub-scripts to run, although the standard sequence represents the optimal configuration for most users. All setup scripts and sub-scripts include ‘-help’ functions describing their operations, and are extensively commented to facilitate modifications. Following execution, a fully functional setup is generated in approximately 15 minutes. For users interested in prototyping new devices, the framework provides template Arduino code files as a starting point.

The Arduino template code supports rapid prototyping and includes the following sections:

- Arduino IDE and Library Installation – Instructions to configure the development environment.
- Configure Pins – Setup and initialization of inputs and outputs: Inputs: analog and digital reads (within the main loop), I2C, and Serial inputs. Outputs: digitalWrite, PWM, and Serial prints (within the main loop).
- Control Constants, Variables and Functions - Initialisation of control variables and functions, including support for control strategies such as Bang-Bang, PID, and timed control.
- Wi-Fi Setup and Initialization - WPA-Enterprise configuration or WPA-Personal configuration
- MQTT (TLS) Setup and Initialisation – Configuration of MQTT communication, including callback functions, publishing, and subscription with defined actions.
- Webserver Setup and Initialisation – Web interface template with on/off buttons and input fields.
- void setup() Function – Utilises the initialisation functions defined above.
- void loop() Function – Executes the main code process.

This structured approach ensures that users can quickly deploy the framework while providing a flexible foundation for device prototyping and adaptation.

### Augmenting a Heating Mantle to Produce a Turbidostat Bioreactor

In addition to the generation of novel devices, the LAS3R framework enables rapid and low cost augmentation of existing hardware in the lab to support automation. This easy and low cost way to accelerate research can significantly reduce manual repetitive tasks. In our example, we modify a heating mantle (common laboratory device used to uniformly heat round bottomed flasks with stirring) with inbuilt temperature control, and turn it into a turbidostat bioreactor through addition of an inline optical density (OD) sensor along with culture media influx and waste efflux pumps. The turbidostat enables maintenance of microbial culture density (measured by OD) at a setpoint in bulk culture of up to 1 L. This is very useful for biological protocols where a specific cell density is needed, or during continuous directed evolution experiments. With a total add-on cost of less than £20 per device (+ raspberry pi node £100) this augmentation is far cheaper than the purchase of commercial bioreactors, which can cost upwards of £1000 for those with equivalent functionality and capable of handling similar volumes. The utility of our setup was demonstrated by generating growth curves for E. coli MG1655 and implementing a controlled growth regime involving repeated dilutions within defined bounds to calculate accurate growth rates (Fig 5). The doubling time of our strain at 37 °C was calculated as (31.96 +-0.26 min). Details of the augmented bioreactor setup, along with a fully integrated bioreactor featuring controllable heating, cooling, and stirring, are described at github.com/KaviHShah/LAS3R Bioreactor.

**Fig 5.**
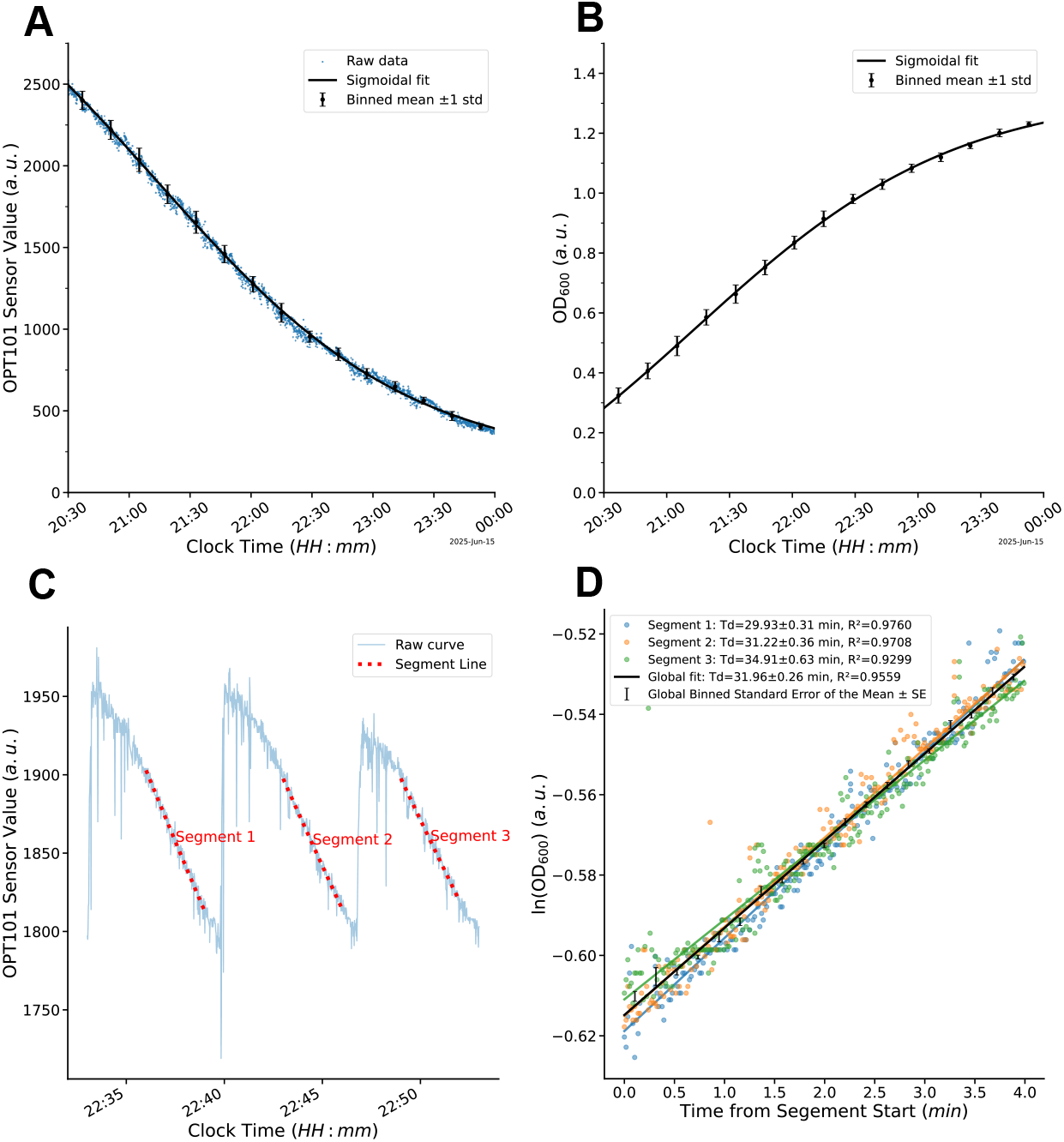
Bioreactor growth curves and growth rate calculation. **A:** Raw Sensor Signal Attenuation - Photodiode data (digital counts, a.u.) showing the decrease in light transmission as E. coli MG1655 density increases during growth. Data is fitted with an exponential decay model. **B:** OD_600_ Curve - Raw digital counts from (A) transformed using a calibration curve (S1 Fig 1) against reference spectrophotometer readings to provide standardized OD_600_ values. **C:** Repeated Dilution Regime - Time-course data where culture is diluted repeatedly when raw readings fall below 1800 a.u. to maintain exponential growth; specific segments (labelled red) are extracted for growth rate analysis. **D:** Log-Linear Growth Analysis - Natural log of calibrated OD_600_ values (*ln*[*OD*_600_]) with linear regressions utilized to determine specific growth rates. Individual replicates (coloured) and global fits are plotted. Mean doubling time: 31.96±0.26 min.

### Template Script Assisted Light Controller Prototyping

To demonstrate the ease of prototyping utilising the template script, a light controller box was created to keep light intensity at a set point. This could be used for example in a plant growth chamber. It took approximately two hours to prototype the light controller. During this period, the ESP32 microcontroller script was updated to limit how quickly the LED output PWM duty cycle changes after a new setpoint is applied to improve stability; the PID constants and cumulative error clamping constants were optimised; and the setup and ESP32 code was further modified to include a backup LED which was turned on in the event of failure of the first. This along with addition of a system daemon to periodically reassert setpoint values and modification of the hostapd configuration to enable rapid reconnection, improved the setup further. This prototype was used for subsequent setup robustness testing (below).

### SPOF Analysis Demonstrates Robustness

To demonstrate, evaluate, and improve the robustness of the setup to multiple different potential faults, Single point of failure analysis (SPOF) was conducted utilising the light controller prototype (Fig 6). The ability to handle faults is crucial in laboratory experiments, where their impact must be minimised to preserve experimental integrity. We demonstrate that light controller setup produced using the LAS3R lab automation framework is robust to most major points of failure. In particular, when there is a failure of the Raspberry Pi Network or Communication, the ESP32 microcontroller based regime continues unimpeded or is halted, whichever is the preferred behaviour for the user. When these services resume, data collection and operation resumes as normal. A second highlight is that the setup can handle full restoration of functionality following an unprotected power failure of the Raspberry Pi (simulated by a loss of power without backup). All the required services resume and data collection restarts as intended. Thirdly, the failover LED effectively takes over upon hardware failure. A final highlight is that the data collection and setup can effectively handle mobile microcontroller devices which enter and exit range, connecting and collecting data when the device is in range. Further data with tandem triplicate tests for each failure mode can be found in the supplementary (S1 Fig 2).

**Fig 6.**
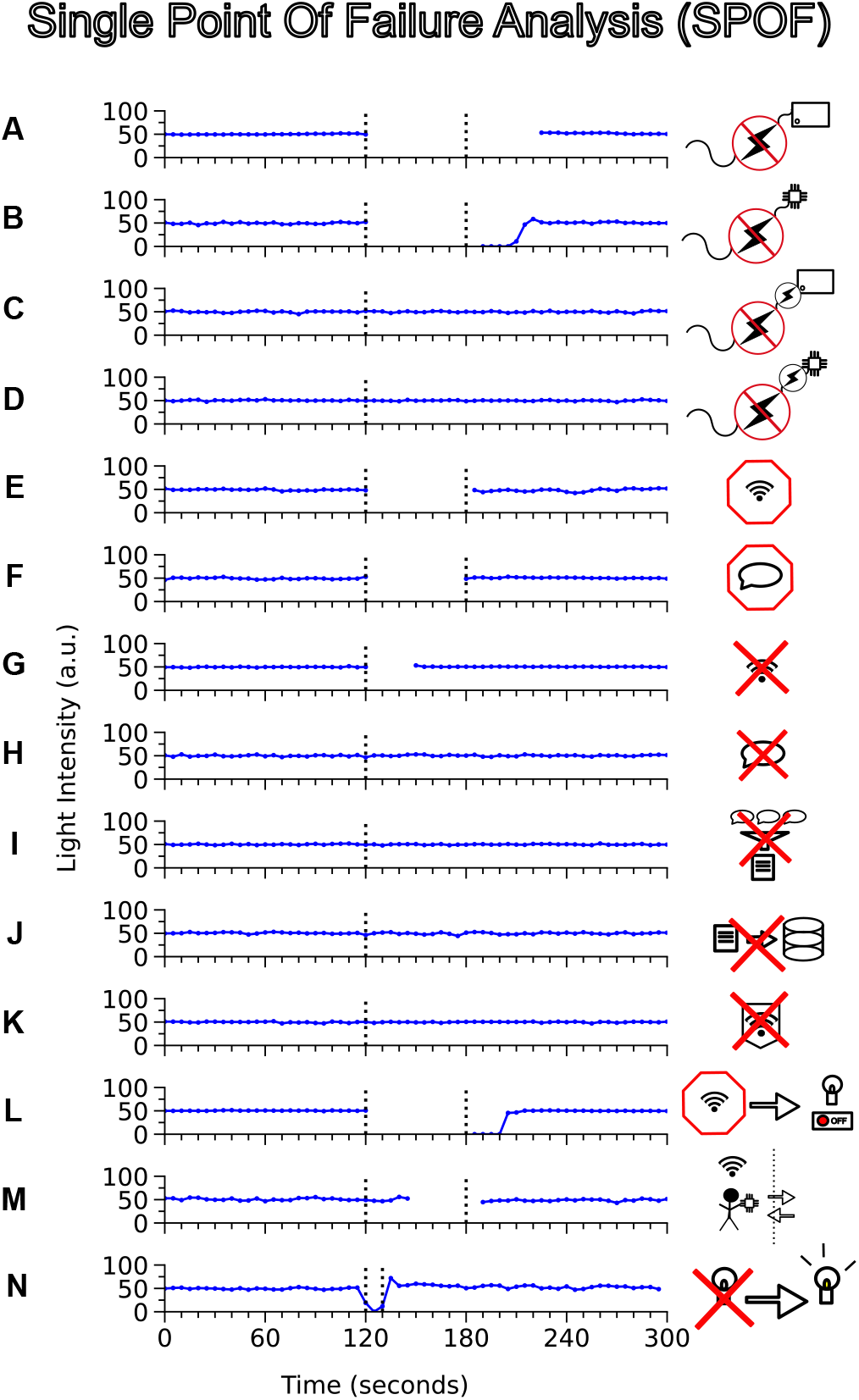
Single Point of Failure (SPOF) analysis demonstrating system robustness and decentralised control. The light intensity PID controller was configured to maintain a steady-state setpoint of 50 arbitrary units (a.u.). Experimental perturbations were initiated at t = 120 s and, where applicable, reversed at t = 180 s. All tests were performed in triplicate to ensure reproducibility (See S1 Fig 2 for expanded data). **A:** Recovery after loss of power to the Raspberry Pi (unbuffered); system services autonomously re-initialized upon reboot and data collection resumed, with the microcontroller maintaining the 50 a.u. setpoint in the interim via local control logic. **B:** Recovery after loss of power to the microcontroller; upon reboot, the ESP32 successfully resynchronized with the Raspberry Pi, transitioning from 0 a.u. back to the 50 a.u. setpoint. **C–D:** Integration of a backup Uninterruptible Power Supply (UPS) for the Raspberry Pi (C) and ESP32 (D), respectively; demonstrated that power failures result in no disruption to service or data collection. **E–F:** Controlled disruption (graceful termination) of network services via the hostapd daemon (E) and communication via the mosquitto broker (F); while data collection was temporarily suspended, the PID loop maintained a constant 50 a.u. intensity. **G:** Non-graceful network termination; abrupt service interruption was followed by autonomous Wi-Fi resumption within 30 s, with light intensity maintained at the target throughout the recovery period. **H–J:** Non-graceful termination of the mosquitto broker (H), telegraf data harvester (I), and influxdb collector (J); these failures resulted in no disruption to light level control and negligible impact on data continuity. **K:** Failure of the freeradius service similarly demonstrated no effect on data collection, as established network connections remained active. **L:** Programmable safety-state behavior; loss of network connectivity triggered an automated shutdown of the light output, with full restoration to 50 a.u. occurring immediately upon reconnection. **M:** Dynamic range and mobility; moving the device out of network range at t = 120 s and returning at t = 180 s demonstrated seamless reconnection and setpoint persistence. **N:** Simulated hardware fault; severing the primary LED circuit initiated an automated failover to a redundant light source with a programmed 10 s latency to facilitate visual verification of the transition.

**Fig 7.**
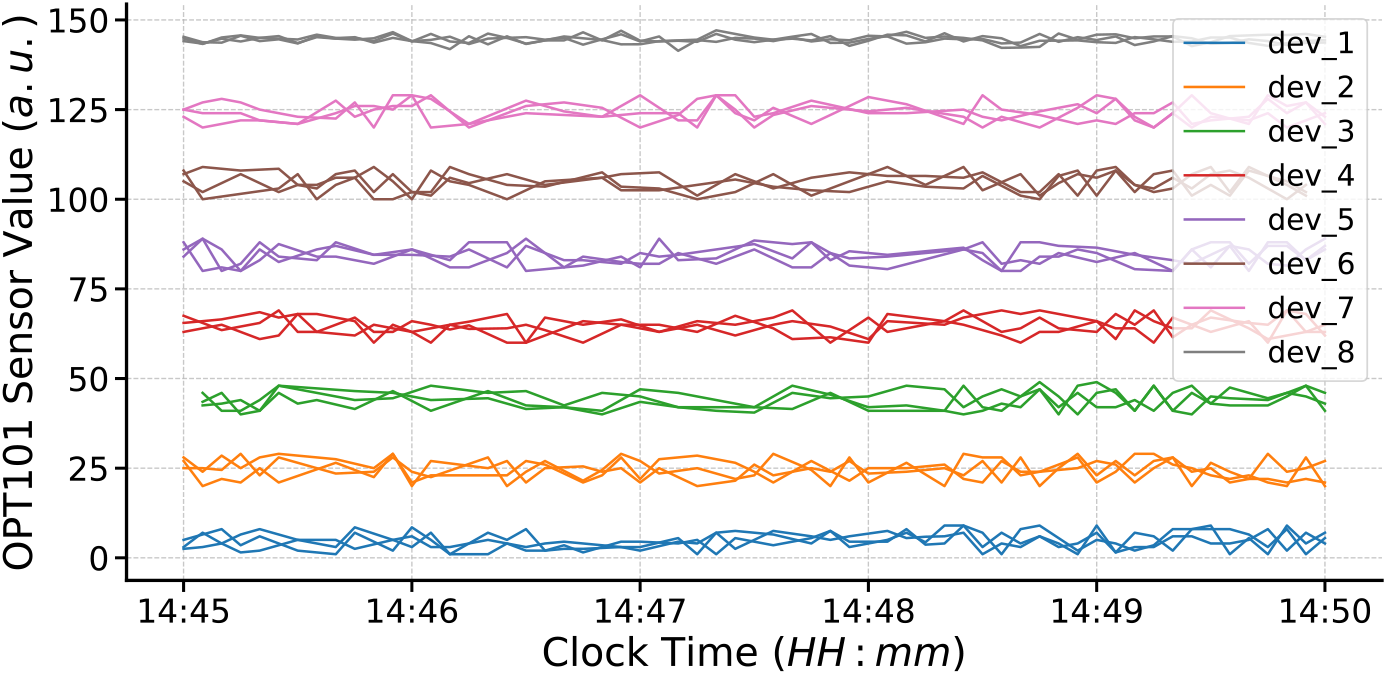
Demonstration of framework scalability. Continuous data streaming from eight simultaneously connected ESP32 devices, each simulating three sensors (24 total). Device set-points were configured at 20 intervals. Sensor readings are binned in 5-second intervals and plotted as single lines, with lines from the same device colored consistently.

### Scalability - Multi Device Management with Raspberry Pi

To assess scalability, we demonstrated simultaneous data streaming from eight independent devices to a single Raspberry Pi each with three sensors (Figure). Although the absolute device limit was not systematically evaluated, we estimate that the firmware would probably support approximately 10–12 connected devices. Whilst three sensors were simulated, the Raspbery Pi and ESP32s can support many more inputs and outputs. The system is further scalable through the integration of multiple Raspberry Pis, for example by distributing services such as message brokering or FreeRADIUS authentication across separate Raspberry Pi’s, or by migrating to enterprise-grade hardware capable of supporting a larger number of concurrent connections. As the setup is built on Debian Linux, it can be readily ported to other hardware platforms running a Linux operating system with minimal modification. Importantly, this architecture is highly cost-effective when using Raspberry Pi’s and ESP32 microcontrollers: a single Raspberry Pi 5 costs approximately £100, while each ESP32-based device costs less than £10, in addition to modest peripheral hardware expenses.

### Code and Documentation

The aim of this project is to use low-cost, open-source software and hardware, to enable easy adoption, and allow users to modify the LAS3R framework to suit their needs. To this end, documentation including a full tutorial documenting device setup, common errors and points of failure, and troubleshooting tips, is provided for users. Security considerations and modifications for more stringent environments are also discussed in the documentation. Furthermore, alternative software and modifications, to support alteration of the setup for different purposes are also discussed. For users wishing to deploy the LAS3R framework, the documentation is the place to start: kavihshah.github.io/LAS3R.

## Conclusion

The presented LAS3R lab automation framework provides a simple, secure, scalable and robust integrated solution for laboratory automation, enabling biologists and users with minimal electronics expertise to rapidly prototype and deploy custom devices. Future developments aim to expand the range of devices for which template code is available, and to increase support for diverse data types, including images. Moreover, fostering a collaborative open-source community will facilitate the generation and sharing of code, supporting a broader spectrum of laboratory devices tailored to varied user needs.

## Acknowledgments

This work was supported by the Department of Genetics – Trinity Hall Award, School of Biological Sciences DTP PhD Studentship.

## References

1. Holland I, Davies JA. Automation in the life science research laboratory. Frontiers in bioengineering and biotechnology. 2020;8:571777.

2. Cizauskas C, DeBenedictis E, Kelly P. How the past is shaping the future of life science: The influence of automation and AI on biology. New Biotechnology. 2025;88:1–11.

3. Steel H, Habgood R, Kelly CL, Papachristodoulou A. In situ characterisation and manipulation of biological systems with Chi. Bio. PLoS biology. 2020;18(7):e3000794.

4. Quero FJ, Aidelberg G, Vielfaure H, Huon de Kermadec Y, Cazaux S, Pandi A, et al. qByte: An open-source isothermal fluorimeter for democratizing analysis of nucleic acids, proteins and cells. PLoS Biology. 2025;23(5):e3003199.

5. Mohanan SMP, Russell K, Duncan S, Kiang A, Lochenie C, Duffy E, et al. FluoroPi device with smartprobes: a frugal point-of-care system for fluorescent detection of bacteria from a pre-clinical model of microbial keratitis. Translational vision science & technology. 2023;12(7):1–1.

6. Collins JT, Knapper J, Stirling J, Mduda J, Mkindi C, Mayagaya V, et al. Robotic microscopy for everyone: the OpenFlexure microscope. Biomedical Optics Express. 2020;11(5):2447–60.

7. Gome G, Waksberg J, Grishko A, Wald IY, Zuckerman O. OpenLH: open liquid-handling system for creative experimentation with biology. In: Proceedings of the Thirteenth International Conference on Tangible, Embedded, and Embodied Interaction; 2019. p. 55–64.

8. McDermott S, Kotar J, Collins J, Mancini L, Bowman R, Cicuta P. Using old laboratory equipment with modern Web-of-Things standards: a smart laboratory with LabThings Retro. Royal Society Open Science. 2024;11(8).

9. Laith AE, Jaouni H, Mihyar A. Addressing cyberbiosecurity challenges in the modern era of biotechnology and artificial intelligence: cyberbiosecurity in the age of biotechnology and AI. Global Biosecurity. 2025.

10. Raspberry Pi Foundation. Raspberry Pi 5; 2025. Single-board computer. https://www.raspberrypi.org/products/.

11. Raspberry Pi Foundation. Raspberry Pi OS 64-bit (Debian-based); 2025. Release date: 13 May 2025. https://www.raspberrypi.org/software/operating-systems/.

12. Raspberry Pi Foundation. Raspberry Pi Imager; 2025. Version 1.8.5, used to image Raspberry Pi OS 64-bit. https://www.raspberrypi.org/software/.

13. The OpenSSL Project. OpenSSL: Cryptography and SSL/TLS Toolkit; 2025. Version: 3.0.17 1 Jul 2025. https://www.openssl.org/.

14. Jouni Malinen and the hostapd developers. hostapd: Host Access Point Daemon; 2023. Version 2:2.10-12+deb12u2 (Debian 12 package). https://w1.fi/hostapd/.

15. Simon Kelley and the dnsmasq developers. dnsmasq; 2023. Version 2. 90-4 (Debian package). http://www.thekelleys.org.uk/dnsmasq/doc.html.

16. FreeRADIUS Project. FreeRADIUS; 2022. Version 3.2.1. https://freeradius.org.

17. Eclipse Foundation and the Mosquitto developers. Eclipse Mosquitto; 2022. Version 2.0.11-1.2+deb12u1 (Debian package). https://mosquitto.org/.

18. The Apache Software Foundation. Apache HTTP Server; 2024. Version 2.4.62-1 deb12u2. https://httpd.apache.org/.

19. Canonical Ltd. Uncomplicated Firewall (UFW); 2023. Version 0.36.2. https://launchpad.net/ufw.

20. InfluxData, Inc. Telegraf; 2025. Installed from Debian package via apt-get. https://github.com/influxdata/telegraf.

21. InfluxData, Inc. InfluxDB 2; 2025. Installed from Debian package via apt-get. https://github.com/influxdata/influxdb/tree/main-2.x.

22. Eclipse Foundation, Roger Light, Pierre Fersing and developers. Paho MQTT Python Client Library; 2024. Version 2.1.0 (latest stable), accessed 2026–04-08. https://pypi.org/project/paho-mqtt/.

23. Arduino. Arduino Integrated Development Environment (IDE); 2025. Version 2.3.7 (macOS build). https://www.arduino.cc/en/software.

24. Espressif Systems. ESP32 Board Package for Arduino IDE; 2025. Version 3.1.1. https://github.com/espressif/arduino-esp32.

25. Arduino LLC and WiFi101 developers. WiFi101 Library for Arduino; 2020. Version 0.16.1. https://github.com/arduino-libraries/WiFi101.

26. Arduino-ESP32 Contributors. WiFiClientSecure, NetworkClientSecure (part of Arduino-ESP32 core); 2025. Arduino-ESP32 GitHub repository. https://github.com/espressif/arduino-esp32/tree/master/libraries/NetworkClientSecure.

27. Espressif Systems and the contributors. ESP-IDF esp eap client Wi-Fi Enterprise; 2025. ESP-IDF GitHub repository. https://github.com/espressif/esp-idf/tree/master/components/wpa_supplicant/esp_supplicant/src.

28. O’Leary N. PubSubClient: Arduino MQTT Client Library (v2.8); 2020. MIT License; Arduino MQTT client library v2.8 release. https://github.com/knolleary/pubsubclient/releases/tag/v2.8.

29. DFRobot. Beetle ESP32 (DFR0575) Wi-Fi & Bluetooth Microcontroller; 2026. Compact ESP32-based wearable Wi-Fi and Bluetooth controller; Arduino-compatible, accessed 2026-04-08. https://www.dfrobot.com/product-1798.html.

30. Adafruit Industries. Adafruit HUZZAH32 ESP32 Feather Board; 2025. ESP32-based Feather development board with Wi-Fi and Bluetooth. https://www.adafruit.com/product/3405.

31. Simon D, Aboba B, Hurst R. The EAP-TLS authentication protocol; 2008. https://www.rfc-editor.org/rfc/rfc5216.html.

32. Rescorla E. The Transport Layer Security (TLS) Protocol Version 1.3. Internet Engineering Task Force (IETF); 2018. RFC 8446. Available from: https://www.rfc-editor.org/rfc/rfc8446.html.

33. Hong S, Kang J, Kwon S. Performance comparison of http, https, and mqtt for iot applications. The International Journal of Advanced Smart Convergence. 2023;12(1):9–17.

