## Supplementary Figures for "LAS3R: A simple, secure, scalable, and robust framework for deploying lab automation devices"

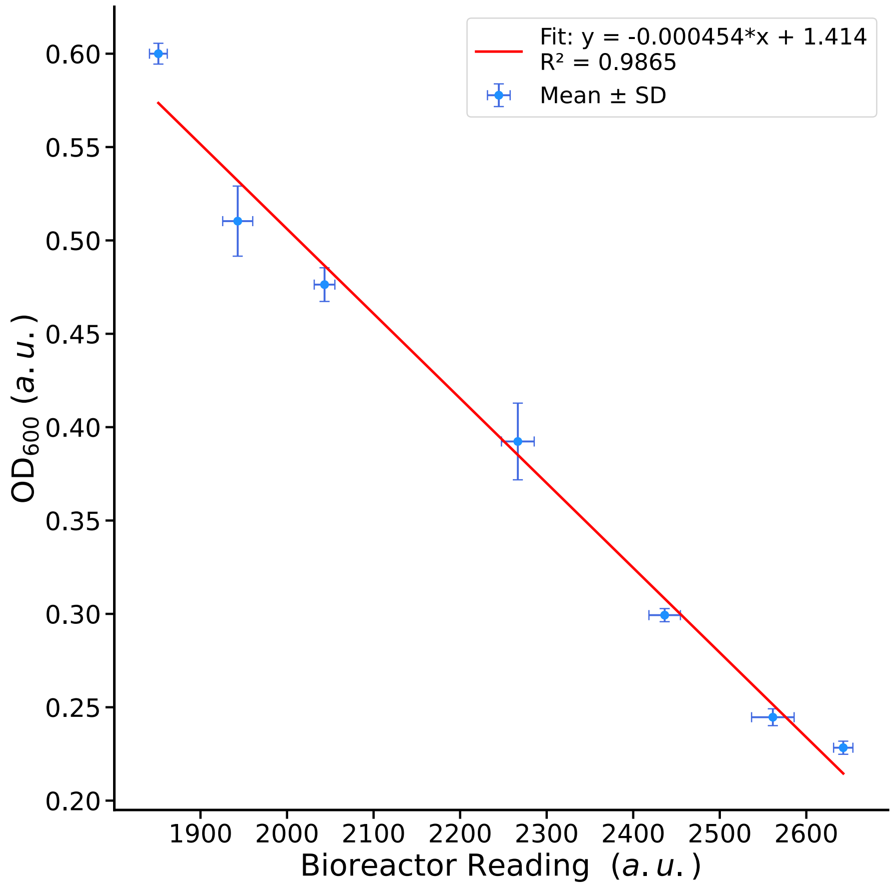


**Fig 1. Bioreactor light sensor vs Spectrophotometer $OD_{600}$.**
Calibration Curve of the Bioreactor OPT101 Light Sensor readings against the GeneQuant 1300 Spectrophotometer enabling of retrieval of $OD_{600}$ values from experiments

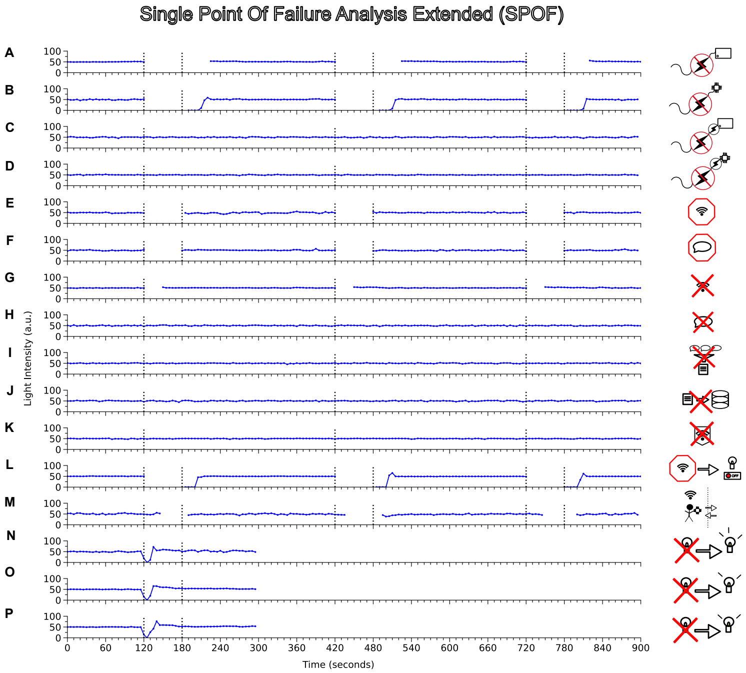

Single Point Of Failure Analysis Extended (SPOF)
A
B
C
D
E
F
G
H
I
J
K
L
M
N
O
P
Light Intensity (a.u.)
Time (seconds)

**Fig 2. Extended single-point-of-failure analysis**
The light intensity PID controller was configured to maintain a steady-state setpoint of 50 arbitrary units (a.u.). Three sequential 300 s cycles were conducted in tandem, yielding a total experimental duration of 900 s per perturbation. Within each cycle, the perturbations were introduced at t = 120 s and, where applicable, reversed at t = 180 s.
**A:** Recovery after loss of power to the Raspberry Pi (unbuffered); system services autonomously re-initialized upon reboot and data collection resumed, with the microcontroller maintaining the 50 a.u. setpoint in the interim via local control logic.
**B:** Recovery after loss of power to the microcontroller; upon reboot, the ESP32 successfully resynchronized with the Raspberry Pi, transitioning from 0 a.u. back to the 50 a.u. setpoint.
**C-D:** Integration of a backup Uninterruptible Power Supply (UPS) for the Raspberry Pi (C) and ESP32 (D), respectively; demonstrated that power failures result in no disruption to service or data collection.
**E–F:** Controlled disruption (graceful termination) of network services via the hostapd daemon (E) and communication via the mosquitto broker (F); while data collection was temporarily suspended, the PID loop maintained a constant 50 a.u. intensity.
**G:** Non-graceful network termination; abrupt service interruption was followed by autonomous WiFi resumption within 30 s, with light intensity maintained at the target throughout the recovery period.
**H–J:** Non-graceful termination of the mosquitto broker (H), telegraf data harvester (I), and influxdb collector (J); these failures resulted in no disruption to light level control and negligible impact on data continuity.
**K:** Failure of the freeradius service similarly demonstrated no effect on data collection, as established network connections remained active.
**L:** Programmable safety-state behavior; loss of network connectivity triggered an automated shutdown of the light output, with full restoration to 50 a.u. occurring immediately upon reconnection.
**M:** Dynamic range and mobility; moving the device out of network range at t = 120 s and returning at t = 180 s demonstrated seamless reconnection and setpoint persistence.
**N-P:** Simulated hardware fault; severing the primary LED circuit initiated an automated failover to a redundant light source with a programmed 10 s latency to facilitate visual verification of the transition. Three independent runs were carried out with time in between to reconnect the primary LED.
